# Dysregulation of Neuroprotective Lipoxin Pathway in Astrocytes in Response to Cytokines and Ocular Hypertension

**DOI:** 10.1101/2023.06.22.546157

**Authors:** Shruthi Karnam, Shubham Maurya, Elainna Ng, Amodini Choudhary, Arzin Thobani, John G Flanagan, Karsten Gronert

**Affiliations:** Herbert Wertheim School of Optometry and Vision Science, University of California Berkeley, Berkeley, California, United States; Infectious Disease and Immunity Program, Herbert Wertheim School of Optometry and Vision Science, University of California Berkeley, CA, United States

**Keywords:** Glaucoma, astrocyte reactivity, lipoxygenase, lipoxin B_4_, neurodegeneration, retina, optic nerve

## Abstract

Glaucoma leads to vision loss due to retinal ganglion cell death. Astrocyte reactivity contributes to neurodegeneration. Our recent study found that lipoxin B_4_ (LXB_4_), produced by retinal astrocytes, has direct neuroprotective actions on retinal ganglion cells. In this study, we aimed to investigate how the autacoid LXB_4_ influences astrocyte activity in the retina under inflammatory cytokine-induced activation and during ocular hypertension. The protective activity of LXB_4_ was investigated *in vivo* using the mouse silicone-oil model of chronic ocular hypertension (n=40). By employing a range of analytical techniques, including bulk RNA-seq, RNAscope in-situ hybridization, qPCR, and lipidomic analyses, we discovered the formation of lipoxins and expression of the lipoxin pathway in rodents (including the retina and optic nerve), primates (optic nerve), and human brain astrocytes, indicating the presence of this neuroprotective pathway across various species. Findings in the mouse retina identified significant dysregulation of the lipoxin pathway in response to chronic ocular hypertension, leading to an increase in 5- lipoxygenase (5-LOX) activity and a decrease in 15-LOX activity. This dysregulation was coincident with a marked upregulation of astrocyte reactivity. Reactive human brain astrocytes also showed a significant increase in 5-LOX. Treatment with LXB_4_ amplified the lipoxin biosynthetic pathway by restoring and amplifying the generation of another member of the lipoxin family, LXA_4_, and mitigated astrocyte reactivity in mouse retinas and human brain astrocytes. In conclusion, the lipoxin pathway is functionally expressed in rodents, primates, and human astrocytes, and is a resident neuroprotective pathway that is downregulated in reactive astrocytes. Novel cellular targets for LXB_4_’s neuroprotective action are inhibition of astrocyte reactivity and restoration of lipoxin generation. Amplifying the lipoxin pathway is a potential target to disrupt or prevent astrocyte reactivity in neurodegenerative diseases, including retinal ganglion cell death in glaucoma.

## Introduction

Glaucoma is a progressive neurodegenerative disease characterized by the degeneration of retinal ganglion cells (RGCs) and their axons [34, 44]. It is a leading cause of irreversible blindness worldwide, and it is estimated that the number of people affected by glaucoma will increase to 112 million worldwide by 2040 [24]. Although intraocular pressure (IOP) is the only modifiable risk factor, most glaucoma patients experience continued but reduced progression. Proposed reasons for this continued degeneration include noncompliance with medication, insufficient IOP reduction, and non-IOP-related causes [6, 15, 21]. There is an unresolved need to develop neuroprotective strategies to prevent or delay the death of RGCs, independent of IOP, in glaucoma [20].

Lipoxins (LXA_4_ and LXB_4_) are eicosanoids generated from arachidonic acid. Their canonical protective actions in various pathological processes include regulating the initial response of innate and adaptive immune cells and promoting the resolution of acute inflammation [32, 38, 52]. We recently identified a noncanonical protective action of lipoxins, namely their direct regulation of RGCs [30]. The human and mouse receptor for LXA_4_, formyl peptide receptor 2 (FPR2/ALX) [39], is expressed in RGCs. A receptor for LXB_4_, the lipoxin with more potent neuroprotective actions in the retina [30], has yet to be identified. The biosynthesis of lipoxins requires the coordinated interaction of 5-lipoxygenase (LOX) with 12/15-LOX enzymes [13, 49] (An overview of lipoxin biosynthesis is included in supplementary Fig. 1). Primary astrocytes generate lipoxins *in vitro* and acute exocytotoxic stress downregulates lipoxin formation and lipoxygenase expression in the retina [30]. However, the regulation of the lipoxin pathway in ocular hypertension has yet to be investigated. More importantly, it is unclear whether retinal astrocyte reactivity, a primary response to ocular hypertension, regulates the astrocyte lipoxin pathway and whether lipoxin formation is a unique phenotype of mouse retinal astrocytes. These knowledge gaps are of interest as therapeutic amplification of the retinal lipoxin pathway is a potential neuroprotective target.

Astrocytes are glial cells in the central nervous system and retina and play a crucial role in maintaining homeostasis. They provide metabolic substrates, growth factors, and trophic support to neurons and are also involved in synapse formation and plasticity [42]. Astrocytes also work closely with the immune system, as they can detect danger signals, respond to secreted cytokines and chemokines, and activate adaptive immune responses [8]. In rodent models of ocular hypertension, astrocyte reactivity in the retina and optic nerve head is a key feature of the retinal injury response [27, 46]. There is also an initial rise in astrocyte reactivity within the distal optic nerve during the degeneration process [43]. Reactive astrocytes can release proinflammatory molecules and oxidative stress-inducing factors, which initiate or amplify neuroinflammation and neuronal damage [9]. Recently, it has been suggested that microglial release of proinflammatory cytokines can induce the formation of neurotoxic reactive astrocytes in the retina and brain [27].

In this study, we aimed to investigate the regulation and functional expression of lipoxin pathways in the mouse retina, optic nerves of mice and primates, and human astrocytes. We also examined whether the neuroprotective action of the autacoid LXB_4_ includes reducing astrocyte reactivity in response to inflammatory cytokine-induced activation and ocular hypertension.

## Materials and Methods

### Mice

C57BL/6J male mice (Jackson Laboratory, Bar Harbor, ME) were allowed to acclimatize to the housing facility for 2 weeks before experimentation. Mouse studies were performed in compliance with the ARVO Statement for the Use of Animals in Ophthalmic and Vision Research, and all experimental procedures were approved by the Institutional Animal Care and Use Committee (IACUC) at the University of California, Berkeley. Mice were maintained in a pathogen-free vivarium under a 12-hour dark and light cycle with *ad libitum* food and water.

### Macaque optic nerves

The macaque optic nerves used for the lipidomic study were from the eyes of a 9-year and 306- day-old female from the Oregon National Primate Research Center (NPRC). For the RNAseq studies, optic nerves were from a 12-year-old male acquired at UC Berkeley. In both cases, the eyes were collected postmortem for approved studies by different research groups, and the optic nerves were donated to the Gronert/Flanagan Lab as discarded tissue.

### Induction of ocular hypertension by injection of silicone oil

Ocular hypertension was induced in the 8-week-old C57BL/6J mice by injection of pharmaceutical-grade silicon oil (Alcon silikon 1000, Geneva, Switzerland) in the anterior chamber of both eyes, a model of chronic ocular hypertension that was established by Jie Zhang and colleagues [11, 50, 51]. For these experiments the volume of silicone oil was reduced to 1.2 μL to enable a slower onset of mild ocular hypertension induced retinal injury. Briefly, 8-week-old mice were anesthetized by an intraperitoneal injection of ketamine/xylazine (100 mg/kg and 10 mg/kg, respectively). One drop of topical anesthetic (proparacaine hydrochloride, 0.5%, Akorn, Lake Forest, IL) was applied to the cornea of both eyes before the injection. Silicone oil (1.2 μL) was injected into the anterior chamber of both eyes through a 30 G Hamilton glass syringe. After injection, a drop of antibiotic (tobramycin ophthalmic solution 0.3%) was placed on the eye. Sterile PBS (phosphate-buffered saline) was injected into another set of mice and used as a control. Mice were euthanized at 2, 4, and 8-week time points following injections for various experimental procedures.

### IOP Measurements/Pupil Dilation

IOP was monitored once every 2 weeks until 8 weeks after silicon oil injection using the TonoLab tonometer (Colonial Medical Supply, Londonderry, NH). In this model, the pupil needs to be dilated sufficiently to expose it beyond the coverage of the silicone oil droplet, which leads to the reconnection of the front and back chambers of the eye [11]. Each measurement of intraocular pressure (IOP) can be seen as a form of pupil dilation treatment. The dilation of the pupil was accomplished following the protocol outlined in the published literature [11, 50, 51].

### Cell Culture

Cultured primary human cortical astrocytes (HA1800) were purchased from ScienCell (Carlsbad, CA) and treated with the cytokines: IL-1α, TNF- α, and C1q for 24 hours [19]. Astrocytes were treated with LXB_4_ (Cayman Chemicals, Ann Arbor, MI) (1 μM) or vehicle for 20 mins before adding cytokines. After 24 hours, the cells were collected in TRIzol (Invitrogen, Waltham, MA) for RNA isolation or ice-cold methanol for lipidomic analysis. For primary retinal astrocytes and optic nerve astrocytes, four mouse retinas were pooled and digested using papain (Worthington, Lakewood, NJ) and plated on poly-D-lysine coated flasks with astrocyte growth media (Lonza, Basel, Switzerland), changing the media every 3 days. Over 95% of the cultured cells were identified as astrocytes through immunofluorescence staining for GFAP and vimentin and were negative for RGCs, microglia, and oligodendrocyte markers.

### Bulk RNA-seq

Total RNA from macaque optic nerves (n=2) was extracted using TRIzol extraction. Sequencing libraries were prepared using the SMARTer v4 Ultra Low Input RNA Kit (Clontech, Mountain View, CA). Sequencing was performed on NovaSeq S4 150PE platform (Illumina, San Diego, CA). Library preparation and RNA-seq were performed at QB3 Genomics, UC Berkeley, Berkeley, CA, RRID: SCR_022170. A quality check of FASTQ files was performed with FastQC v.0.11.9, and adapter sequences were trimmed using Trim Galore! v.0.6.6. Trimmed reads were mapped on mouse genome version mm39 (University of California Santa Cruz) with the STAR alignment tool v.2.7.1a using RefGene annotation. The total number of read counts mapped for each gene was extracted from BAM files using FeatureCount v.1.5.3. The count matrix was imported into Rstudio v.4.2.0. It was normalized using the Transcripts Per Million method by NormalizeTPM. Mean values of normalized counts for each gene were plotted for visualization.

### Electroretinography

Before ERG, the mice underwent overnight dark adaptation. They were then anesthetized and received tropicamide (1%, Akorn, Lake Forest, IL) and phenylephrine (2.5%, Paragon BioTeck, Portland, OR) to dilate their pupils. The Celeris system (Diagnosys LLC, Lowell, MA) was used to conduct the ERG measurement, with stimulus intensities ranging from -5.90 to 2.25 log cd.s/m^2^. The tail subdermal needle electrode served as ground. A positive scotopic threshold response (pSTR) elicited with an intensity of -2.50 log cd.s/m^2^ was used to represent RGC function (average of 20 repeats, interstimulus interval of 2 seconds).

### Optical Coherence Tomography

Before imaging, mice received anesthesia and were given lubricant eye drops. We used a Bioptigen spectral domain OCT system (Envisu R2300, Durham, NC) for imaging. Our previous studies described the OCT image acquisition and analysis protocols [28, 29]. We captured an en-face retinal fundus image with the optic nerve head in the center, using 1.8 x 1.8 mm rectangular scanning. Each replication involved 100 B-scan images (1536 A-scans for each B-scan). Retinal layer B-scan images were analyzed by masked observers using ImageJ software. We quantified the thickness of the retinal layer at both left and right locations from the center of the optic nerve head in each retinal quadrant. The average data from both locations represented the thickness value.

### Retinal Flat Mount and RGC Quantification

Eyes were removed at 8 weeks and fixed with 4% paraformaldehyde (PFA). Posterior eyecups were incubated overnight in blocking buffer (10% normal donkey serum and 2% PBS-Triton-X- 100). Eyecups were stained with primary antibodies, GFAP (Abcam, Cambridge, UK), and RBPMS (Phosphosolutions, Aurora, CO) (1:1000 dilution each) and were incubated on a shaker for 48-72 hours at 4 °C. After incubation, eyecups were washed with 0.25% PBST and further incubated with either Alexa Fluor 488- or 594- conjugated secondary antibodies (Invitrogen; at 1:1000 dilution) under dark conditions for 2 hours at room temperature. After washing with PBST, the retinas were gently detached, and four perpendicular clover cuts were made.

ProLong™ Gold Antifade Mountant (#P36930, Invitrogen, Eugene, OR) was added to mount the retinas. Retinas were imaged on a Nikon Ti Eclipse microscope using a 20x objective. A minimum of 5-6 independent specimens (biological replicates) were analyzed for each flat mount. There was no observed RGC loss in the central region. To count peripheral RGCs, eight fields were sampled from the peripheral areas of each retina using a 20x lens on a Nikon Ti Eclipse microscope. (Supplementary Figure 2 displays a flat-mounted retina that defines the central and peripheral regions). RGCs were counted using ImageJ software. The percentage of RGC survival was determined by calculating the ratio of surviving RGC numbers in the injured eyes compared to the control eyes.

### Immunohistochemistry

The eyes were fixed in 4% PFA and transferred into 30% sucrose. Tissues were then embedded in OCT media and cut into 10-μm-thick sections using a CM1900 Cryostat (Leica, Wetzlar, Germany). Primary human astrocytes were fixed with 4% PFA for 20 mins. The cells were permeabilized with 0.3% Triton®-X-100 PBS for 5 mins at room temperature. Sections and cells were blocked using a blocking buffer (5% donkey serum in 0.3% Triton X-100 containing PBS) for 30 mins. Primary antibodies against GFAP (Abcam), RBPMS (Phosphosolutions), LCN2 (Invitrogen), 5-LOX (Abcam), Pax2 (Abcam), Vimentin (Invitrogen), and CD44 (Proteintech) were added to blocking buffer and incubated overnight at 4 °C. Sections and cells were washed and incubated with Alexa Fluor 488- and 594- conjugated secondary antibodies (Invitrogen; at 1:1000 dilution) for double labeling for 2 hours at room temperature under dark conditions. They were then washed and mounted using ProLong™ Gold Antifade Mountant, and images were captured using a Nikon Ti Eclipse microscope.

### RNAscope in situ hybridization

RNA in situ hybridization for *Alox5*, *Alox15*, and *Fpr2* mRNA was performed manually using RNAscope Multiplex Fluorescent Reagent Kit v2 Assay (Advanced Cell Diagnostics, Inc., Newark, CA) according to the manufacturer’s instructions. Briefly, 10 μm of PFA fixed frozen retina and optic nerve sections were pretreated with heat and protease (30LJmins, 40 °C) before hybridization with the target probes. Tissue was incubated in primary amplification probes and Opal^TM^ 690 dye fluorophore (2-hour primary probe, 30LJmins AMP1, 30LJmins AMP2, 15LJmins AMP3, and 30 mins fluorophore at 40 °C) and washed between steps with RNAscope washing buffer. Each sample was quality controlled for RNA integrity with an RNAscope® probe specific to PPIB RNA and for background with a probe specific to bacterial dapB RNA. Specific RNA staining signals were identified as red, punctate dots. The tissue was counterstained with DAPI. After mounting in ProLong™ Gold Antifade Mountant, images were acquired on a Nikon Ti Eclipse microscope using a 40x objective. The RNAscope probes used were as follows: *Alox15* (Cat. No: 539781-C1), *Alox5* (Cat. No: 436101-C2), and *Fpr2* (Cat. No: 509281-C3).

### qPCR

RNA was isolated from mouse retinal tissue and human brain astrocytes and quantified according to our published protocol [30]. *Gapdh* was used as the reference gene. The mouse primers used in this study are listed in Table. Primers for the human genes used in this study are listed in Table 2 (please refer to supplementary material for primer details).

### Liquid Chromatography-tandem mass spectrometry (LCDMS/MS)

Mouse retinas and macaque optic nerves were immediately snapped, frozen on dry ice, and stored at −80 °C before processing for lipidomic analyses. Eicosanoids and PUFAs in the cells and conditioned media from astrocyte cultures were identified and quantified by LCLJMS/MS according to our published protocol [30]. Briefly, deuterated internal standards (15-HETE-d_8_, LXA_4_-d_5_, and AA-d_8_) were added to all samples before processing to calculate class-specific recoveries. Frozen tissues were placed in MeOH and processed in a refrigerated beat homogenizer. Supernatants were extracted using C18 solid-phase columns. Extracted lipids were analyzed by LCLJMS/MS using an AB SCIEX 4500 QTRAP mass spectrometer. Analysis was carried out in negative ion mode, and eicosanoids and PUFAs were quantitated using scheduled multiple reaction monitoring (MRM) using from 3 to 4 specific transition ions for each analyte. The peak identification and integration criterion was a signal-to-noise ratio above 5:1. For quantification, calibration curves and HPLC retention times for each analyte were established with synthetic standards (Cayman Chemicals).

### Statistical Analysis

All values are presented as the mean ± standard error of the mean (SEM) of n observations, where n represents the number of animals studied or independent *in vitro* experiments. Statistical analysis was performed using GraphPad Prism 9.0 (GraphPad Software, San Diego, CA). One-way ANOVA followed by post hoc Tukey’s multiple comparison test or unpaired Student’s t-test was used to compare intergroup differences. The results were considered statistically significant when P<0.05.

## Results

### The lipoxin pathway is expressed in mouse retina, optic nerve, and retinal astrocytes

The expression of lipoxin pathways was investigated in the retina, optic nerve, and retinal astrocytes of C57BL/6J mice. The genes in question are known to be expressed at levels that cannot be detected using single-cell transcriptomics, a known limitation of this method.

Additionally, since there are no specific antibodies for detecting lipoxin pathways in mouse retina and optic nerve, we used RNAscope and qPCR for our investigations. RNAscope in situ hybridization was performed on the retina and optic nerve region cross-sections to determine the cell-specific expression of *Alox5, Alox15, and Fpr2*. The data showed that the lipoxin circuits were localized in the ganglion cell layer (GCL), inner nuclear layer (INL), and outer nuclear layer (ONL) of the retina, and in the optic nerve region (Fig. 1A). The quality of the RNA was confirmed using positive and negative controls, as illustrated in Supplementary Figure 3. To confirm the expression of the lipoxin circuits in retinal astrocytes, qPCR was performed on cultured primary mouse retinal astrocytes. The results showed that *Alox5, Alox15, and Fpr2* were expressed with average Ct values of 28.23±0.3, 29.07±0.3, and 28.58±0.2, respectively (n=3).

**Figure 1.**
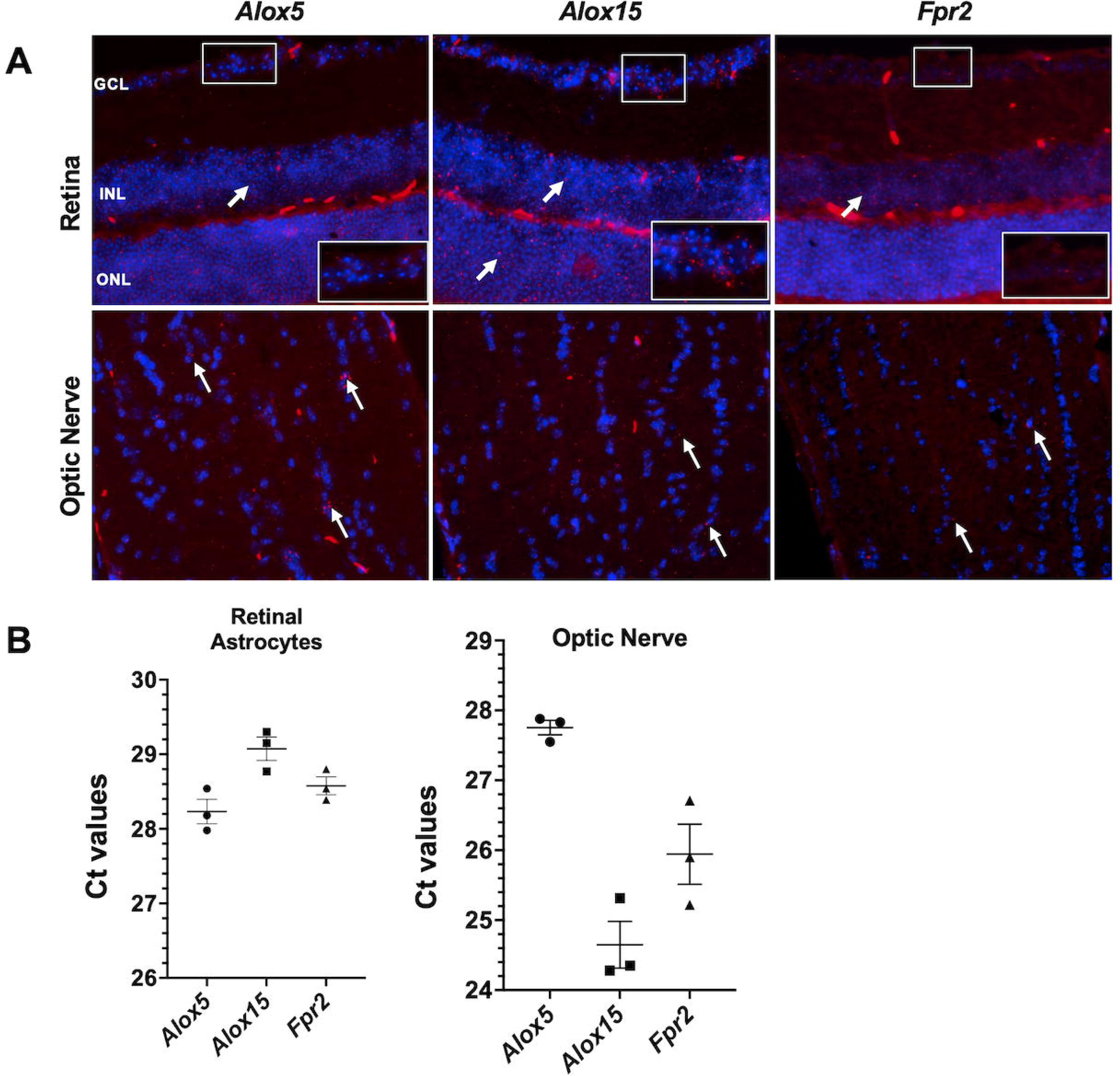
Expression of lipoxin pathway in mouse retina and optic nerve. (A) Representative images of RNAscope ISH (red) (indicated with white arrows) of retina and optic nerve using mRNA probes for *Alox5*, *Alox15*, and *Fpr2*. Nuclei were labeled with DAPI to distinguish nuclear layers of the retina. All images were obtained using a 40x objective. Scale bar = 50 µm. (n=3). Labels: ganglion cell layer (GCL), inner nuclear layer (INL), outer nuclear layer (ONL). **(B)** Bar graph represents Ct values of lipoxin pathway genes in retinal astrocytes and optic nerves analyzed by qPCR (n=3).

Notably, higher RNA expression was observed in the optic nerves with average Ct values of 27.75±0.2 (Alox5), 24.65±0.6 (Alox15), and 25.94±0.7 (Fpr2) (n=3) (Fig. 1B). These results confirm expression of the lipoxin pathway in the retina and optic nerve of mice, including retinal astrocytes.

### Lipoxin pathway is expressed in macaque optic nerves and human primary brain astrocytes

To explore expression of the lipoxin pathway in primates, we conducted RNA-seq analyses on macaque optic nerves. Results established high expression of *ALOX15*, moderate expression of *ALOX5* and *ALOX5AP*, and lower expression of *ALOX12, ALOX15B, FPR2,* and *FPR3* genes (Fig. 2A). Consistent with gene expression, lipidomic analysis confirmed active eicosanoid pathways and lipoxin formation in primate optic nerves as evidenced by the presence of released arachidonic acid and significant levels of 5-HETE, 12-HETE, 15-HETE and LXB_4_ (Fig. 2B).

**Figure 2.**
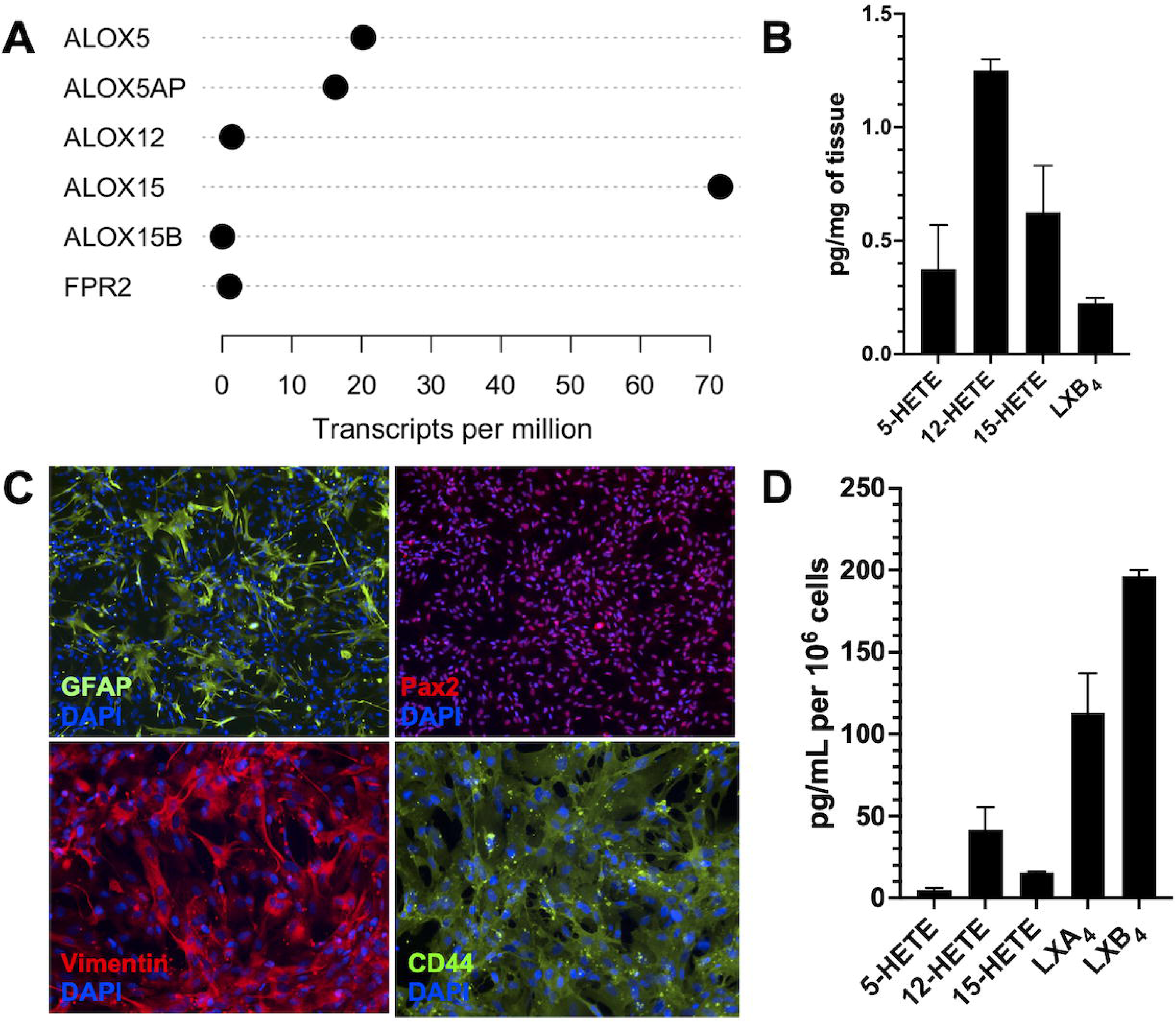
Expression of lipoxin pathway in optic nerve of primates and primary human brain astrocytes. (A) RNA-seq of the macaque optic nerve showing lipoxin pathway genes (n=2). **(B)** LC/MS/MS-based lipidomic quantification of endogenous lipid mediators in primate optic nerve (n=2). **(C)** Representative immunofluorescence images of primary human brain astrocytes stained with GFAP, Pax2, Vimentin, and CD44. **(D)** LC−MS/MS-based lipidomic quantification of endogenous lipid mediators in human brain astrocytes (n=2).

The cell identity of the primary human brain astrocyte line was validated by positive staining with astrocyte-specific markers GFAP, Vimentin, Pax2, and CD44 (Fig. 2C). Primary human astrocytes, like primate optic nerves, also expressed the lipoxin biosynthetic pathway, namely *ALOX5* and *ALOX15,* and the LXA_4_ receptor *FPR2*, and lipidomic analyses confirmed endogenous formation of eicosanoids and lipoxins. Interestingly, cultured human brain astrocytes generated both LXA_4_ and LXB_4_ (Fig. 2D). These data demonstrate that neuroprotective lipoxins are formed within non-human primate optic nerves and can be generated by primary human brain astrocytes.

### Validation of the mild chronic model of ocular hypertension

IOP and RGC density changes were assessed in control and silicone oil-injected eyes of mice. Fig. 3A shows that IOP was significantly elevated in the silicone oil-injected eyes compared to control eyes. After 8 weeks mice were euthanized, and RGC density was assessed by staining flat-mounted retinas with the RGC marker, RBPMS (Fig. 3B). Peripheral RGC loss of 17% was observed in silicon oil-injected eyes compared to control eyes (Fig. 3C). Changes in RGC function were measured longitudinally using ERG measurements of pSTR. The average waveforms of pSTR were elicited by a flash intensity of -2.5 log cd.s/m^2^ at 8 weeks (Fig. 3D). The results indicated a loss of RGC function in hypertensive eyes compared to control normotensive eyes. Additionally, loss in ganglion cell complex (GCC) thickness, a clinically relevant biomarker of nerve fiber loss, was measured longitudinally by OCT (Fig. 3E). Mild ocular hypertension caused a significant 10% relative loss (p<0.05) compared to normotensive control eyes (Fig. 3F). Overall, these results confirm that the injection of 1.2 μl of silicone oil established mild, chronic ocular hypertension and mild damage to the RGCs after 8 weeks. The sustained elevated intraocular pressure (IOP) resulted in mild morphological and functional loss of retinal ganglion cells (RGCs).

**Figure 3.**
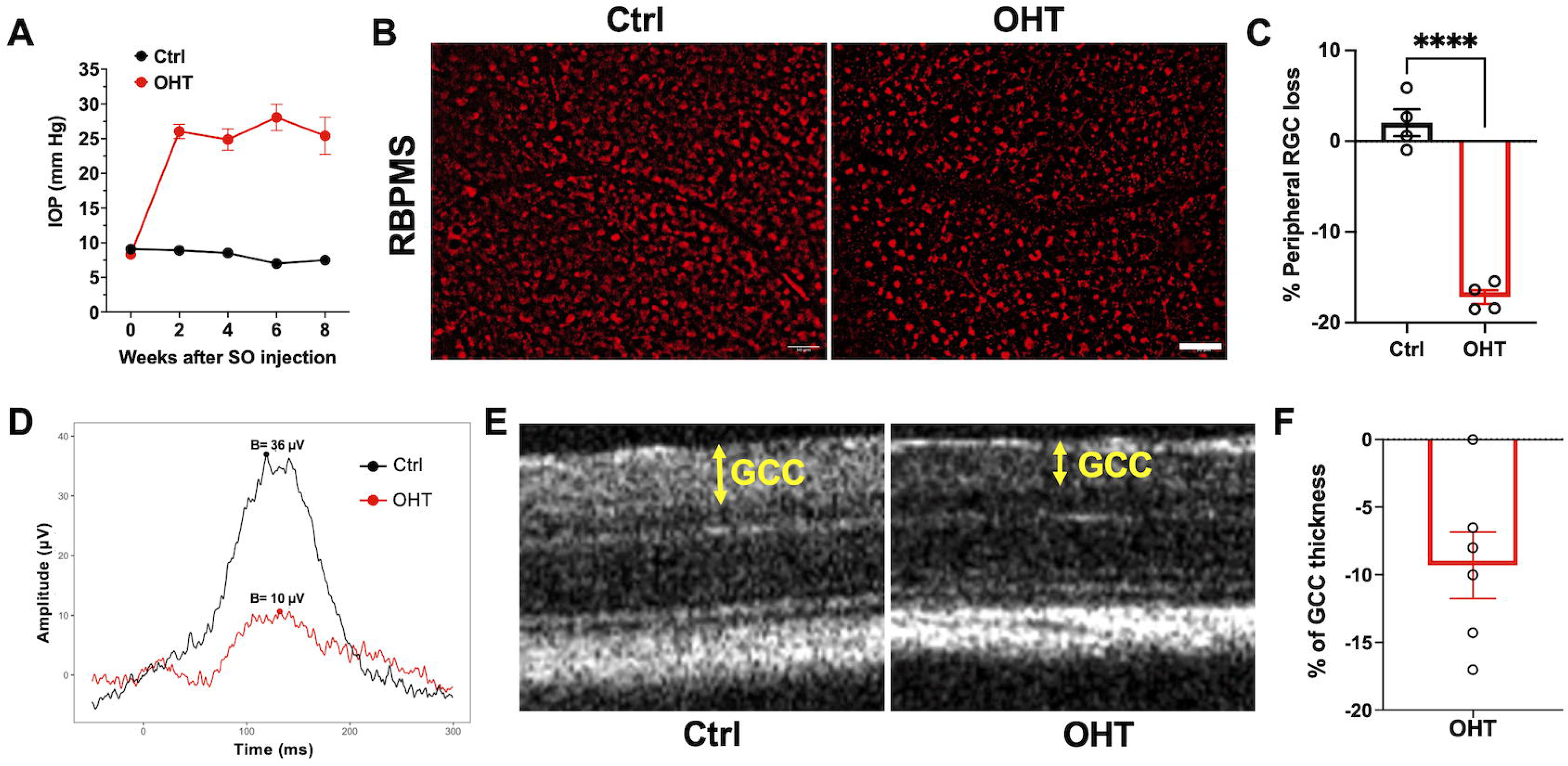
Validation of the silicone oil model. (A) IOPs were increased after silicone oil (SO) injection (n=20). **(B)** Representative confocal images of flat-mounted retina sections showing surviving RBPMS-positive (red) RGCs at 8 weeks. Scale bar = 50 µm. **(C)** Bar graph indicates % of peripheral RGC loss. **(D)** Average ERG waveforms of RGC responses (pSTR, -2.5 log cd.s.m^-2^) at 8 weeks for each group. **(E)** Representative OCT images of mouse retina at 8 weeks. GCC: ganglion cell complex, including RNFL, GCL, and IPL layers, indicated by double-ended yellow arrows. **(F)** Bar graph indicating the % reduction in GCC thickness at 8 weeks. (n=4-6). ****p< 0.001. (Ctrl- Control; OHT- ocular hypertension).

### Dysregulation of lipoxin pathway in response to chronic mild ocular hypertension

Lipoxin pathway gene expression was analyzed in retinas from normotensive and hypertensive eyes at 2, 4, and 8 weeks to investigate the regulation of lipoxin pathway in response to chronic mild ocular hypertension. The results demonstrate that *Alox5AP* (5-lipoxygenase activating protein, FLAP), which is a cofactor required for full 5-LOX enzymatic activity, was significantly upregulated at 2 and 4 weeks, while there was no change in the levels of *Alox5* in response to ocular hypertension. Interestingly, the levels of *Alox15* and *Fpr2* were significantly upregulated initially at 2 and 4 weeks but significantly downregulated at 8 weeks (Fig. 4A). To investigate the functional dysregulation of the lipoxin pathway, four retinas were pooled at 4 weeks and analyzed by LCLJMS/MS-based lipidomics to establish formation and changes in eicosanoid and PUFA profiles. Lipidomic analysis demonstrated that arachidonic acid (AA) and the 15-LOX pathway marker, 15-HETE, were significantly decreased in response to ocular hypertension. In contrast, the 5-LOX pathway markers, 5-HETE, and 4-HDHA were significantly increased (Fig. 4B). These findings indicate that expression of the homeostatic lipoxin pathway is regulated during ocular hypertension and that activity of 5-LOX and 15-LOX is dysregulated.

**Figure 4.**
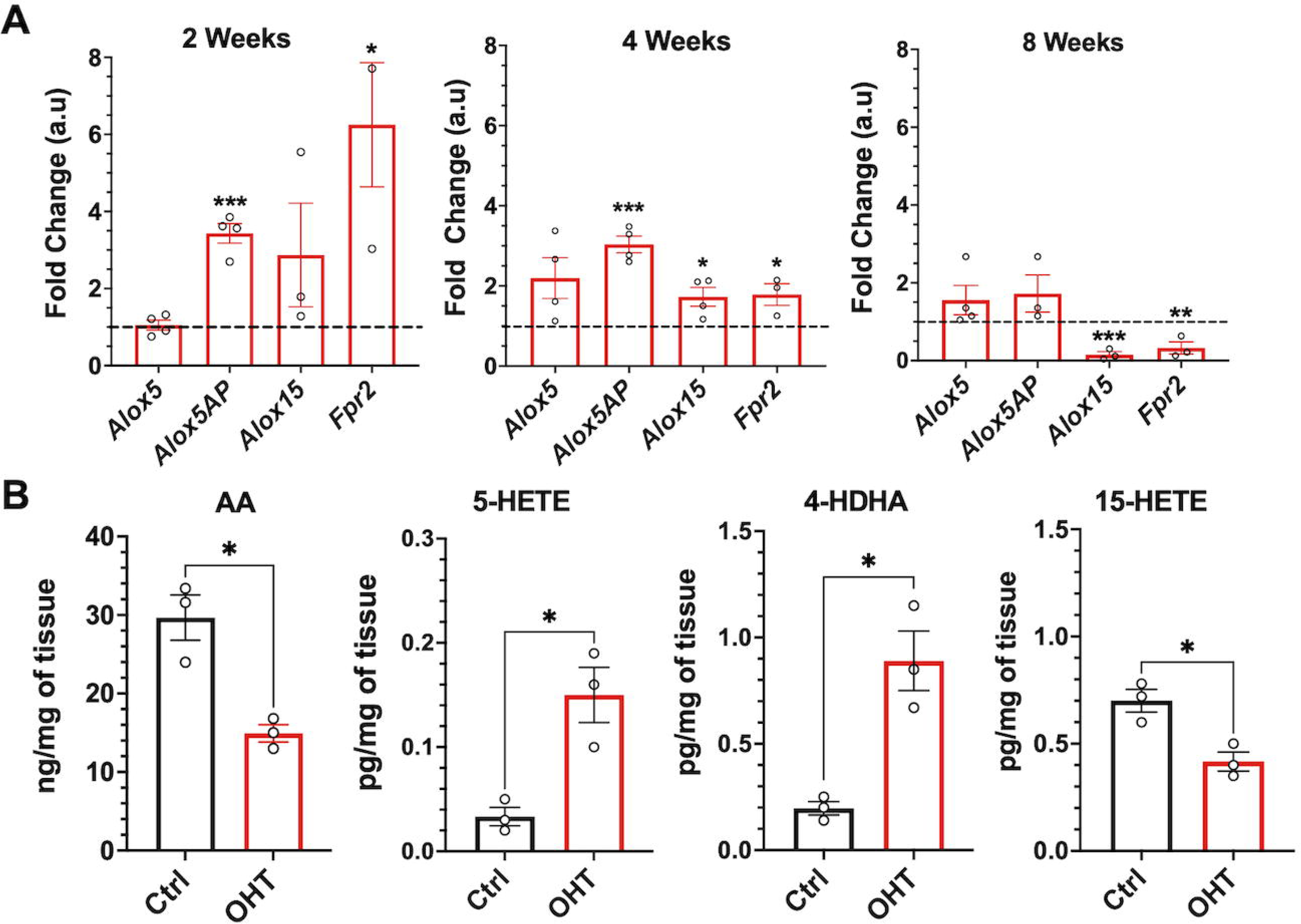
Regulation of lipoxin pathway in response to ocular hypertension. (A) The time course response of lipoxin pathway genes was analyzed by qPCR at 2, 4, and 8 weeks. The results were normalized to GAPDH and age-matched controls. n=3-4 (2 retinas were pooled for each replicate). **(B)** LC/MS/MS-based lipidomic quantification of endogenous lipid mediators in the mouse retina at 4 weeks, n=3 (4 retinas were pooled for each replicate). *p< 0.05, **p<0.01, **p<0.001. (Ctrl- Control; OHT- ocular hypertension).

### Activation of Müller glia and astrocyte reactivity markers in response to ocular hypertension

The impact of mild ocular hypertension on astrocyte and Müller glial reactivity was assessed by immunofluorescence staining of retinal cross-sections with cell-specific reactivity markers from normotensive and hypertensive eyes. Expression of GFAP, an intermediate filament protein that is a marker of macroglial reactivity, was induced and upregulated in Müller glia and astrocytes, as early as 2 weeks after ocular hypertension (Fig. 5A) and remained upregulated in the retina for 8 weeks and was coincident with loss of RGCs as evidenced by RBPMS staining in the GCL layer (Fig. 5B). GFAP staining of retinal flat mounts also established marked reactive astrogliosis after 8 weeks of mild ocular hypertension (Fig. 5C). To investigate macroglia reactivity further, we measured glutamate aspartate transporter (GLAST) gene expression, which is highly expressed in Müller cells as part of their homeostatic function to support RGCs. GLAST expression was significantly decreased after 2 and 4 weeks of ocular hypertension (Fig. 5D).

**Figure 5.**
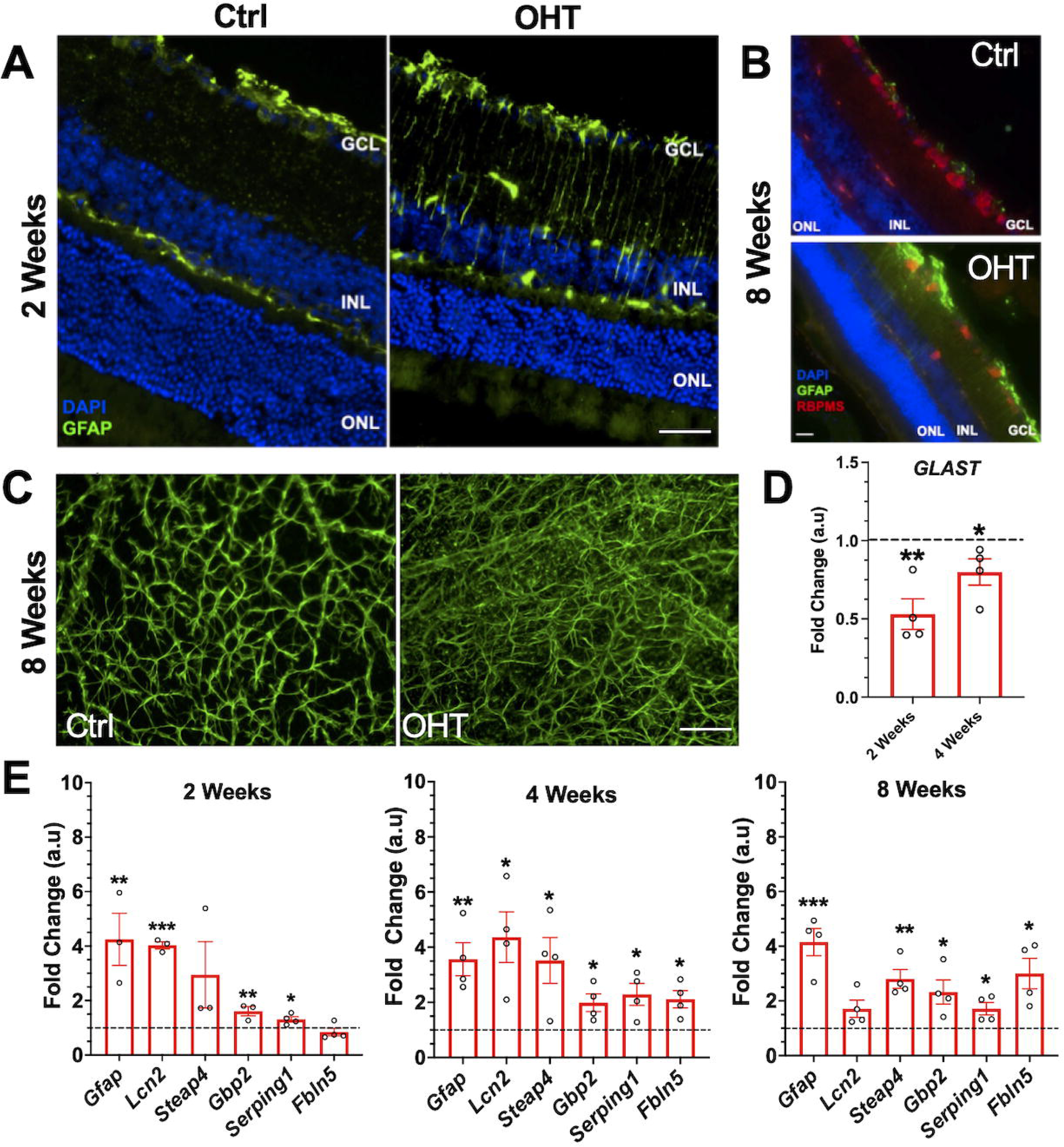
Regulation of astrocyte reactivity in response to ocular hypertension. (A) Representative immunofluorescence images of GFAP (green) at 2 weeks, and **(B)** GFAP (green) RBPMS (red) at 8 weeks of ocular hypertension. Nuclei were labeled with DAPI. All images were obtained using a 40x objective. Scale bar = 50 µm. Labels: ganglion cell layer (GCL), inner nuclear layer (INL), outer nuclear layer (ONL). **(C)** Representative images of a portion of flat-mounted retinas stained with GFAP (green) at 8 weeks. **(D)** Muller glia reactivity gene, GLAST, was analyzed by qPCR at 2 and 4 weeks. The results were normalized to GAPDH. **(E)** Astrocyte markers were significantly upregulated at 2, 4, and 8 weeks when analyzed by qPCR. The results were normalized to GAPDH and controls. n=4 (2 retinas were pooled for each replicate). *p< 0.05, **p<0.01, **p<0.001. (Ctrl- Control; OHT- ocular hypertension).

This dysregulation has also been previously observed in rat models of optic nerve injury [33]. Gene expression of signature astrocyte reactivity markers (*Gfap, Lcn2, Steap4, Gbp2, Serping1, and Fbln5*) were significantly upregulated in response to ocular hypertension at 2, 4, and 8 weeks compared to that in normotensive controls (Fig. 5E). These findings confirm that the early occurrence of Müller glia and astrocyte reactivity is a characteristic feature in this mouse model of mild ocular hypertension-induced retinal injury.

### Therapeutic LXB_4_ treatment reduces astrocyte and Muller glial reactivity in the mouse retina

We next assessed whether the neuroprotective actions of lipoxins in ocular hypertension [30] include regulation of macroglia reactivity. Therapeutic LXB_4_ methyl ester treatment (n=8, 5 ng/g IP; 0.5 ng/g topical) was initiated after 1 week of ocular hypertension and administered every other day until the 4th week. The dosing and preparation of lipoxins were based on previous studies [5, 30]. qPCR analysis of markers of macroglia reactivity established that treatment with LXB_4_ resulted in a significant upregulation of GLAST. In addition, LXB_4_ treatment significantly downregulated astrocyte reactivity markers such as *Lcn2* and *Steap4*. The levels of *Gbp2*, *Serping1, and Fbln5* remained unchanged (Fig. 6). These findings identify a potentially new mechanism for the neuroprotective actions of LXB_4_ treatment, namely reversing macroglia reactivity and restoring their homeostatic function.

**Figure 6.**
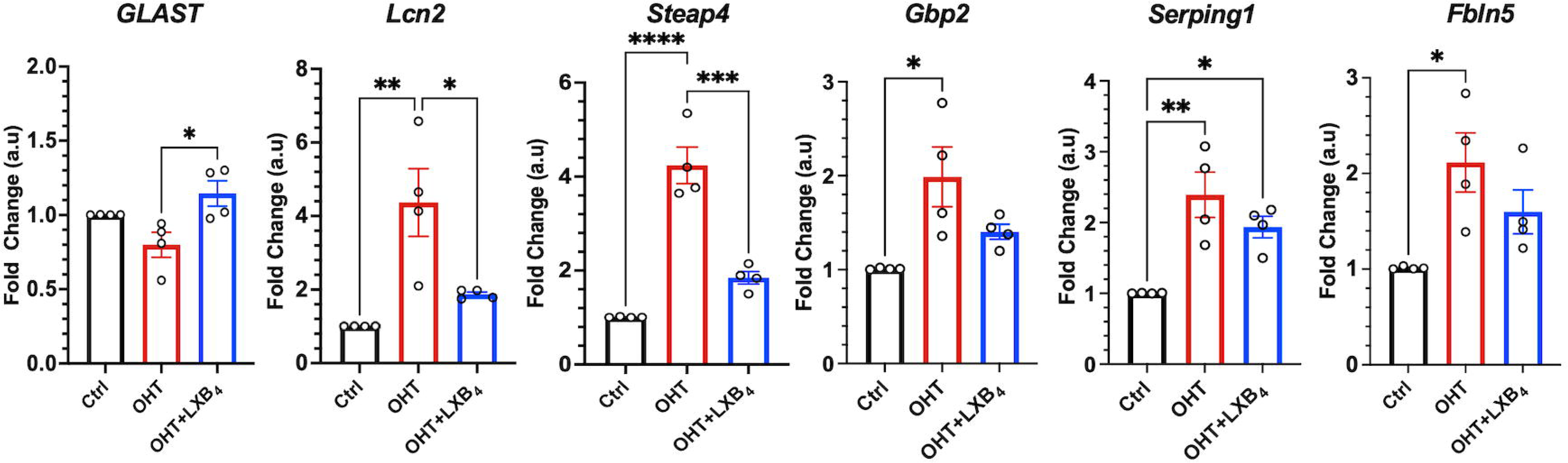
Effect of LXB_4_ treatment on astrocyte reactivity markers in the mouse retina. LXB_4_ methyl ester treatment (n=8, 5 ng/g IP; 0.5 ng/g topical) was started after one week of silicone oil injection every other day until the 4^th^ week. Treatment with LXB_4_ modulates astrocyte reactivity genes in the ocular hypertension model, n=4 (2 retinas were pooled for each replicate). (Ctrl- Control; OHT- ocular hypertension). *p< 0.05, **p<0.01, ***p<0.001.

### LXB_4_ treatment modulates the lipoxin pathway in primary human brain astrocytes

To establish macroglia-specific actions for LXB_4_, we used primary human brain astrocytes that express a fully functional lipoxin pathway identical to that found in the mouse primary retinal astrocytes (Fig. 2D). To induce astrocyte reactivity, cells were treated with established inflammatory cytokines, which are also generated in the retina in response to ocular hypertension [19]. qPCR analysis confirmed that cytokine treatment significantly upregulated reactive astrocyte markers, including *LCN2, SERPING1,* and *C3*, in primary human brain astrocytes. LXB_4_ treatment significantly downregulated these astrocyte reactivity markers (Fig. 7A). Immunofluorescence analysis of LCN2 was consistent with the qPCR findings. These results establish direct regulation of astrocyte reactivity as a new bioaction of LXB_4_.

**Figure 7.**
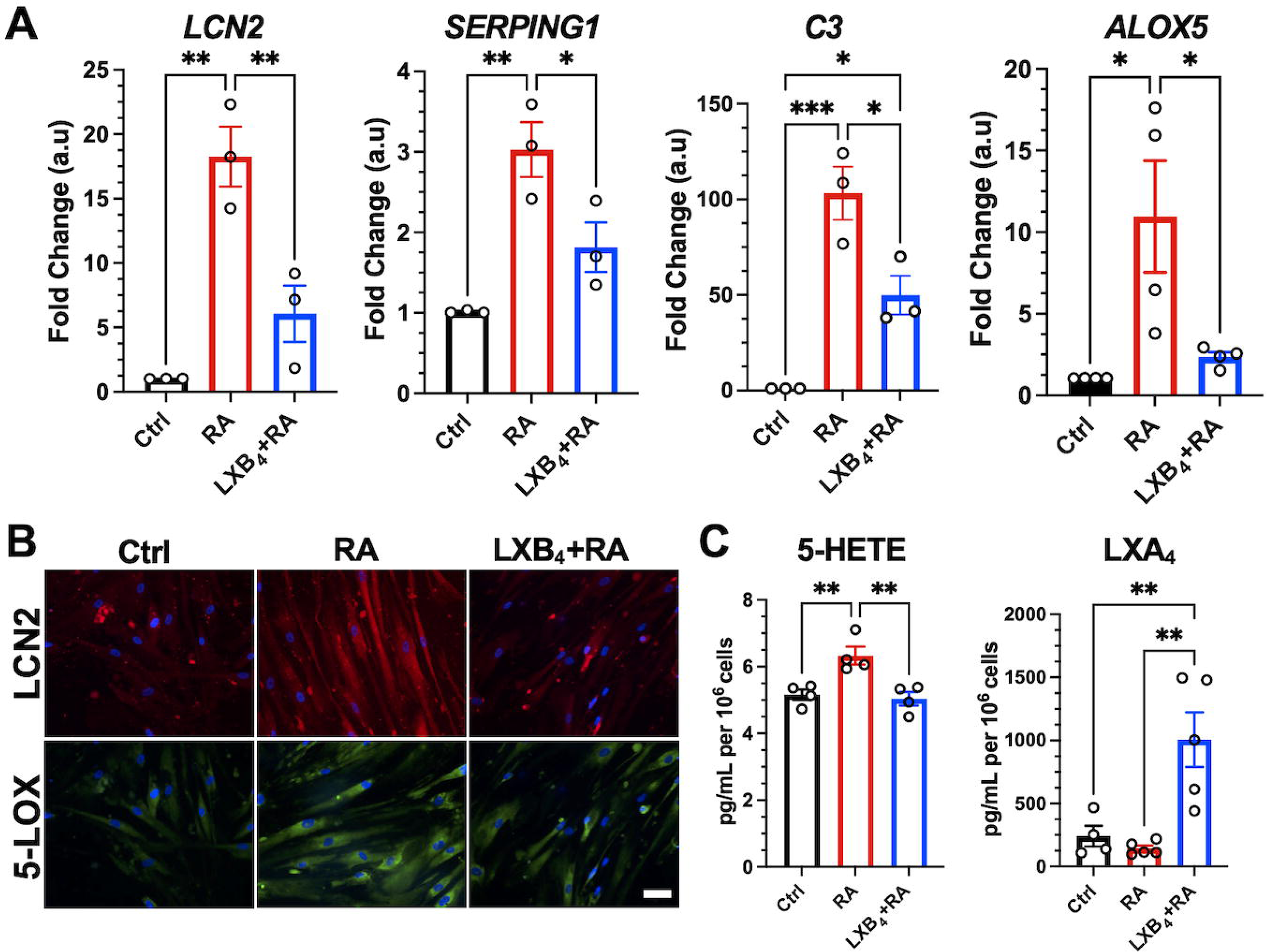
Effect of LXB_4_ treatment on lipoxin pathway in human brain astrocytes. (A) Astrocyte reactivity genes analyzed by qPCR. The results were normalized to GAPDH. Data are presented as the mean±SEM, n=3-4. **(B)** Representative immunofluorescence images of LCN2 and 5-LOX. Nuclei were labeled with DAPI. **(C)** LCLJMS/MS-based lipidomic quantification of endogenous lipid mediators, n=4-5. (Ctrl-untreated control astrocytes, RA- reactive astrocytes). *p< 0.05, **p<0.01, ***p<0.001.

Reactive astrocytes also showed increased gene expression of 5-LOX (ALOX5) compared to controls. LXB_4_ treatment reduced 5-LOX expression in reactive astrocytes (Figs. 7A and B). Hence, we examined the functional consequence of LXB_4_ regulation of 5-LOX expression in reactive astrocytes. Consistent with changes in 5-LOX gene expression (Fig 7B), 5-HETE formation was significantly increased in reactive astrocytes, and LXB_4_ treatment reduced 5-HETE generation. More importantly, reactive astrocytes generated less LXA_4_, but LXB_4_ treatment restored and amplified LXA_4_ formation in reactive astrocytes (Fig. 7C).

Concentrations of added LXB_4_ in media remained high after 24 hours, so we were unable to assess whether LXB_4_ treatment also amplified endogenous astrocyte LXB_4_ formation. These findings suggest LXB_4_ restores the homeostatic phenotype, and both recovers and amplifies neuroprotective lipoxin formation in cytokine-challenged primary human astrocytes.

## Discussion

Glaucoma results in the gradual loss of retinal ganglion cells (RGCs), leading to irreversible vision loss [18]. Astrocyte reactivity is a key pathological mechanism that contributes to the neurodegeneration of RGCs in glaucoma, as astrocytes play a crucial role in maintaining the health and function of RGCs [2]. Recently, we identified that lipoxins (LXA_4_ and LXB_4_) derived from astrocytes promote neuroprotection against acute and chronic injury in the retina by directly acting on RGCs [30]. However, the regulation of lipoxin formation in the retina and the extent of its neuroprotective activity in glaucoma remains to be defined.

Our study revealed that astrocytes express 5-LOX and 15-LOX, two enzymes essential for lipoxin generation by a single cell [13, 48, 49]. We assessed the functional activity of the lipoxin pathway through LCLJMS/MS-based lipidomics and confirmed the presence of endogenous lipoxin formation and activity of 5-LOX and 15-LOX in rodent and primate astrocytes, retina, and optic nerves. Notably, we observed lipoxin formation in the optic nerves of nonhuman primates, which, to our knowledge, has not been previously reported. Additionally, we demonstrated formation of both LXA_4_ and LXB_4_ by primary human brain astrocytes, aligning with a recent study reporting lipoxin formation in astrocytes of human brain organoids using ELISA [35]. Astrocytes, the most abundant cell type in the central nervous system, play vital regulatory roles in neurogenesis, synaptogenesis, bloodLJbrain barrier permeability, and extracellular homeostasis, impacting various aspects of brain function [41]. These results emphasize the potential significance of the resident neuroprotective lipoxin pathway not only in the retina but also in the brain and optic nerve [25, 30].

5-LOX is required to generate leukotrienes, which are proinflammatory lipid mediators, and this enzyme has been implicated in promoting proinflammatory lipid mediator synthesis in various neurodegenerative diseases [36]. However, 5-LOX is also required for generating lipoxins and most of the proresolving lipid mediators (SPMs) [25, 38, 49]. Hence, the role of 5- LOX is context- and cell-dependent. In the retina, 5-LOX has been identified as the essential enzyme for the protective actions of dietary DHA supplementation in retinopathy of prematurity [37]. Moreover, in 5-LOX knockout mice, autoimmune uveitis pathogenesis is markedly amplified, and in a retinal model of glutamate excitotoxicity, 5-LOX inhibition amplifies retinal ganglion cell death [30]. These findings indicate that in the retina, homeostatic 5-LOX is protective.

The function of 5-LOX, namely the products this pathway generates, depends on the balanced interaction with 15-LOX, an interaction that leads to lipoxin formation. 15-LOX is highly expressed in epithelial cells of healthy mucosal tissues, retinal pigmented epithelial cells, tissue-resident homeostatic macrophages, immune regulatory macrophages and polymorphonuclear leukocytes (PMNs) [3, 14, 16, 47]. 15-LOX has immunoregulatory and wound healing roles important for maintaining tissue homeostasis, especially in the eye [3, 14, 17, 47]. A key feature of an immune-regulatory and pro-resolving phenotype of PMN is high expression of 15-LOX to enable generation of lipoxins [14, 26]. Dysregulation of the balanced expression and function of 5- and 15-lipoxygenases has been implicated in several neurodegenerative diseases [25] and deletion of either 5-LOX or 15-LOX amplifies experimental allergic encephalomyelitis in mice [10]. Our findings reveal a shift in the homeostatic balance of 5-LOX and 15-LOX activities in the retina and astrocytes as a feature of reactivity and ocular hypertension-induced pathogenesis. Astrocyte reactivity and retinal response to ocular hypertension correlated with an increase in 5-LOX and a decrease in 15-LOX activity.

Interestingly, our findings of dysregulated lipoxygenase activity are consistent with previous studies that have reported increased levels of 5-LOX activity in Alzheimer’s disease (AD) [7, 12, 22]. Additionally, a separate study reported decreased levels of 15-LOX in the CA1 region of the hippocampus in patients with moderate to severe stages of AD [31]. In contrast, several studies have demonstrated that inhibition of mouse 12/15-LOX enzymes can reduce ischemic injury and diabetic pathogenesis [1, 45]. In short, there is a complex interplay between 5- and 15- LOX, and their regulation and dysregulation are a key feature of neurodegenerative diseases such as Alzheimer’s disease and glaucoma.

Our study identified a novel target for the neuroprotective effects of LXB_4_, demonstrating its ability to inhibit astrocyte reactivity both *in vitro* and *in vivo*. Astrocytes play a crucial role in maintaining the health and function of RGCs and the vasculature in the retina and optic nerve.

Astrocyte reactivity has been implicated as a key pathological feature in various neurodegenerative diseases, including glaucoma [4, 23]. Our findings establish, for the first time, a direct action of LXB_4_ on macroglia by downregulating key reactive markers in primary human astrocytes *in vitro*. This finding highlights the significance of LXB_4_ as an autocrine signal in astrocytes, similar to its established role in the PMN, the primary cell type that generates lipoxins, but also the primary cell target to downregulate their proinflammatory function during acute inflammation [14, 40, 49]. Thus, our results suggest that LXB_4_ serves as an autocrine signal in astrocytes, downregulating their reactive phenotype.

Previous studies have demonstrated the anti-inflammatory effects of lipoxins in various neuroinflammatory diseases, indicating their therapeutic potential [52]. However, the source of lipoxins in the central nervous system and the potential targets for their protective actions have not been thoroughly investigated. Our study reveals that LXB_4_ treatment restores and significantly amplifies the generation of LXA_4_, another lipoxin with distinct bioactions, in reactive astrocytes. The specific receptor for LXB_4_, distinct from the LXA_4_ receptor Fpr2/ALX, remains to be identified, as does the mechanism underlying the upregulation of lipoxin formation in astrocytes. LXB_4_ treatment reduced amplified 5-LOX activity in reactive human astrocytes, suggesting that restoring the balanced interaction between 5-LOX and 15-LOX may partly explain the restoration of lipoxin production in astrocytes.

## Conclusions

The lipoxin pathway is functionally expressed in retina and brain astrocytes, and optic nerves of rodents and primates, and is a resident neuroprotective pathway that is downregulated in reactive astrocytes. Novel cellular targets for LXB_4_’s neuroprotective action is inhibition of astrocyte reactivity and restoration of lipoxin generation. Amplifying the lipoxin pathway is a potential target to disrupt or prevent astrocyte reactivity in neurodegenerative diseases.

## Abbreviations

LXB_4_: Lipoxin B_4_
LXA_4_: Lipoxin A_4_
IOP: Intraocular pressure
RGCs: Retinal ganglion cells
LOX: Lipoxygenase
ERG: Electroretinogram
OCT: Optical coherence tomography
LC: MS/MS-liquid chromatography-tandem mass spectrometry
AA: Arachidonic acid
PUFA: Polyunsaturated fatty acids
HPLC: High-performance liquid chromatography
ONL: Outer nuclear layer
INL: Inner nuclear layer
GCL: Ganglion cell layer
GCC: Ganglion cell complex
SPM: Specialized pro-resolving lipid mediators
PMN: Poly morphonuclear leukocytes.

## Declarations

### Ethics approval and consent to participate

Animal studies were performed in compliance with the ARVO Statement for the Use of Animals in Ophthalmic and Vision Research, and all experimental procedures were approved by the Institutional Animal Care and Use Committee (IACUC) at the University of California, Berkeley.

### Consent for publication

Not applicable

### Availability of data and material

The RNA sequencing data from the macaque study conducted in this research has been deposited in the GEO under the accession code GSE241131.

## Acknowledgments

We thank Dr. Teresa Puthussery (UC Berkeley), Dr. Joni D. Wallis (UC Berkeley) and the tissue distribution program at Oregon National Primate Research Center (Grant number: P51OD011092) for providing post-mortem macaque optic nerve samples.

## Authors contribution

SK: Conceptualization, Methodology, Validation, Investigation, Formal analysis, Data Curation, Writing- Original Draft, Visualization; SM: Methodology, Validation, Investigation, Formal analysis, Data Curation; EN, AC, AT: performed lipidomic analyses; JF, KG: Conceptualization, Supervision, Resources, Writing-Review & Editing, Funding Acquisition. All authors read and approved the final manuscript.

