## Supplementary Table 1 for "Dysregulation of Neuroprotective Lipoxin Pathway in Astrocytes in Response to Cytokines and Ocular Hypertension"

| **Primers** | **Forward Sequence 5'- 3'** | **Reverse Sequence 5'- 3'** |
| --- | --- | --- |
| *Gapdh* | TGGCCTTCCGTGTTCCTAC | GAGTTGCTGTTGAAGTCGCA |
| *Alox5* | ACAGCTTATCTGCGAGTATGG | GGGAAACACAGGGAGGAATAG |
| *Alox5AP* | GCATGAAAGCAAGGCGCATAA | GGTACGCATCTACGCAGTTCT |
| *Alox15* | GCGACGCTGCCCAATCCTAATC | ATATGGCCACGCTGTTTTCTACC |
| *Fpr2* | GCCAGGACTTTCGTGAGAGAT | GTAGAACTGGTGCTTGAATCACT |
| *Serping-1* | GAACTTGGACCAGGACGCAG | GCTGGTAGCTTCGGGATCTG |
| *Gfap* | GAAACCAACCTGAGGCTGGA | CCACATCCATCTCCACGTGG |
| *GLAST* | GTCGCGGTGATAATGTGGTA | AATCTTCCCTGCGATCAAGA |
| *Steap-4* | CCTCGCCGCAGTAATGGAG | CCCGAAATCTCCTGTTCCGA |
| *Fbln5* | GGCTGTGAGACAAGCAGAAGT | GGTAACAGTGAGTATCATGCGTC |
| *LCN2* | CCACCACGGACTACAACCAG | AGCTCCTTGGTTCTTCCATACA |
| *Gpb2* | GGAGAGTGCTGTGCTGACTT | GGCTACCAGCAGTTCCCTA |

**Table 1: Sequence of Mouse Forward and Reverse Primers**
