## Supplementary Table 2 for "Dysregulation of Neuroprotective Lipoxin Pathway in Astrocytes in Response to Cytokines and Ocular Hypertension"

**Table 2: Sequence of Human Forward and Reverse Prime**

| **Primers** | **Forward Sequence 5'- 3'** | **Reverse Sequence 5'- 3'** |
| --- | --- | --- |
| *GAPDH* | GGAGCGAGATCCCTCCAAAAT | GGCTGTTGTCATACTTCTCATGG |
| *ALOX5* | TCAACTTCGCCAGTACGAC | TCTGCTCAATGGTCACCACG |
| *ALOX15* | TCAGGTTCCCTTGTTACCGC | GTTTCCCCACCGGTACAACT |
| *FPR2* | AGTCTGCTGGCTACACTGTTC | TGGTAATGTGGCCGTGAAGA |
| *SERPING-1* | GGGATGCTTTGGTAGATTTCTCC | GAGGATGCTCTCCAGGTTTGT |
| *LCN2* | GAAGTGTGACTACTGGATCAGA | ACCACTCGGACGAGGTAACT |
| *C3* | GGGGAGTCCCATGTACTCTATC | GGAAGTCGTGGACAGTAACAG |
