## Supplementary Figs for "Dysregulation of Neuroprotective Lipoxin Pathway in Astrocytes in Response to Cytokines and Ocular Hypertension"

**Supplementary Figure 1: An Overview of Lipoxins Biosynthesis**

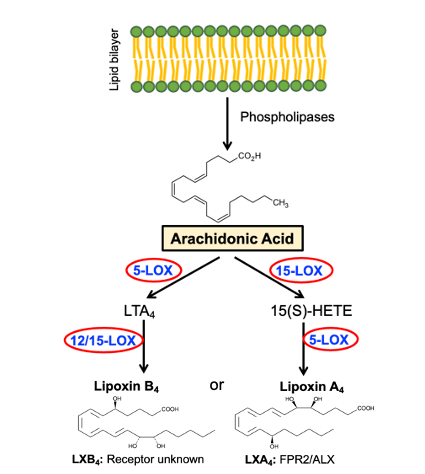

**Suppl. Fig. 1:** LXA_4_ and LXB_4_ are synthesized from arachidonic acid through sequential oxygenation by actions of 5-LOX and 12/15-LOX enzymes. LOX-Lipoxygenases.

**Supplementary Figure 2: Sketch of a flat-mounted retina showing central (C) and peripheral (P) regions.**

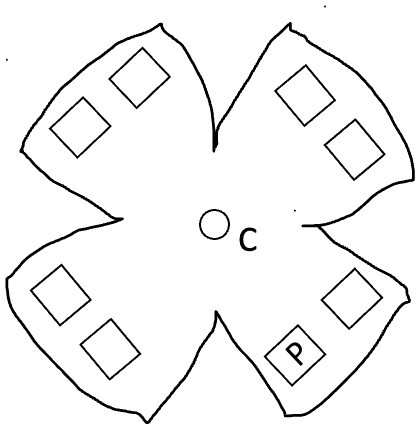

**Suppl. Fig. 2:** After immunostaining, whole retinal flat mounts with RBPMS, eight fields were sampled from each retina's peripheral (indicated with square boxes) areas to determine RGC counts. The circle in the center is the position at which the optic nerve exits the eye.

**Supplementary Figure 3: Confirmation of RNA quality**

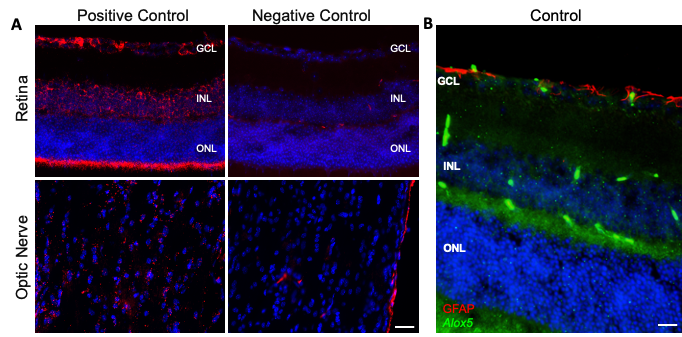

**Suppl. Fig. 3:** A. Using mRNA probes, representative images of positive and negative controls RNAscope ISH (red) of the retina and optic nerve. B. Representative images of RNAscope ISH (green), colocalized with GFAP protein (red) of the retina using mRNA probes for *Alox5*. Nuclei were labeled with DAPI to distinguish the nuclear layers of the retina. All images were obtained using a 40X objective. Scale bar = 50 µm. Labels: ganglion cell layer (GCL), inner nuclear layer (INL), outer nuclear layer (ONL).
